# Taxonomy - for Computers

**DOI:** 10.1101/022145

**Authors:** Nico M. Franz, Beckett W. Sterner

## Abstract

We explore solutions for identifying and reconciling taxonomic concepts that take advantage of the powers of computational representation and reasoning without compromising the suitability of the Linnaean system of nomenclature for human communication. Using the model of the semiotic triangle, we show that taxonomic names must variously achieve reference to nomenclatural types, taxonomic concepts (human-created theories of taxonomic identities), and taxa (evolutionary entities in nature). Expansion of the reference models into temporally transitioning systems shows that the elements of each triangle, and provenance among elements across triangles, are only identifiable if taxonomic names and concepts are precisely contextualized. The Codes of nomenclature, by mandating identifier (name) reuse but not requiring concept-specific identifier granularity, leave the challenge of framing and aligning the symbol/reference instances to human communicators who have superior cognitive abilities in this regard. Computers, in turn, can process greater volumes of narrowly framed and logically aligned data. Comparative, taxonomically referenced biological data are becoming increasingly networked and reutilized in analyses that expand their original context of generation. If we expect our virtual comparative information environments to provide logically enabled taxonomic concept provenance services, then we must improve the syntax and semantics of human taxonomy making – for computers.

---

Much of our understanding of biology is comparative in nature. Communication of comparative information frequently requires biologists to refer to taxonomic entities. This forum paper will suggest that opportunities now being facilitated through computational representations of taxonomic knowledge should motivate comparative biologists to improve the identification and integration of such knowledge. Translation of the semantics produced by taxonomy – herein broadly defined to entail phylogenetics – into the language of computational logic may facilitate a new Semantic Biology where the powers of computational infrastructures are exploited to address complex comparative questions (Deans et al. 2015). We show here that this vision requires careful analysis of the semiotics of human taxonomy making. In particular, we identify elements in contemporary nomenclatural and taxonomic practice that seem to rely so much on uniquely human abilities to contextualize taxonomic information as to impede effective computational processing at larger scales. We explore solutions for taxonomy that take advantage of the powers of computational reasoning without compromising the high degree of suitability of the Linnaean system – as currently implemented – for human communication. These explorations are neither comprehensive nor final, but they point to new and potentially widely impacting pathways for integrating philosophy of language, computational logic, and taxonomic practice. Above all, we hope to encourage comparative biologists to ponder what it would mean to contextualize perceived taxonomic entities – for computers.

## Types, taxa, and concepts – three kinds of taxonomic reference services

Taxonomic names are central to human communication about taxonomic entities (Patterson et al. 2010). To begin understanding why computational logic might struggle with contextualizing taxonomic information, is it helpful to review the kinds of reference services that taxonomic names routinely perform for biologists. We can fathom at least three kinds of services, each with additional subordinate derivations.

The first suite of reference services pertains to the nomenclatural realm. Example statements include: *Andropogon virginicus* is typified by the specimen LINN No. 1211.12. *Andropogon* is typified by *Andropogon virginicus. Andropogon virginicum* is an incorrect spelling of *Andropogon virginicus. Andropogon leucostachyus* Kunth 1816: 187 is a heterotypic, junior synonym of *Andropogon virginicus* Linnaeus 1753: 1046. In the preceding statements, taxonomic names offer communication services by virtue of referring to nomenclatural types. Witteveen (2015a, 2015b) shows that the reference relationship of names and types is one of necessity, and therefore the type method in taxonomy conforms to the Kripke–Putnam causal theory of reference. Dubois (2005: 381–382) applies the term *onomatophores* to type specimens in particular, stating that they “establish an objective link between the real world of organisms and the world of language (nomenclature)”. In short, one critical suite of reference services that taxonomic names provide is manifested in their rigid designation of types.

The second suite of reference services is perhaps most frequently used by the majority of comparative biologists. Example statements include: *Andropogon virginicus* – broomsedge – is widely distributed in the Southeastern United States, and the species produces abundant seeds. *Andropogon virginicus* differs from *Andropogon glomeratus* by having shorter postflowering peduncles. Here the names are used to represent historically, evolutionarily coherent sets of organisms in nature, i.e., taxa.

The third suite includes example statements of the following form. The taxonomic history of *Andropogon virginicus* is full of disagreements. Radford, Ahles and Bell (1968) define *Andropogon virginicus* more broadly than Hitchcock and Chase (1950). In these phrases the taxonomic names most immediately identify taxonomic accounts published by human authors, or perhaps only perceptions of those accounts from the perspectives of other humans. The accounts may have multiple explicit and implicit components (Franz et al. 2015a). Each constitutes an empirically informed theory of the identity and definitional boundaries of the perceived taxon. Henceforth we will refer to these theories as taxonomic concepts (Franz et al. 2008).

We are of the view that all three kinds of reference services – i.e., names referencing types, taxa, and concepts – are very much needed to facilitate effective communication in comparative biology. However, we suggest that the prevailing practice of calling on these services relies on uniquely human abilities for contextualizing name usages, and consequently hampers computational representations of taxonomic content. Below, we describe logic-amenable contingencies that exist between the three different service suites for particular communication needs. To do so we first introduce the semiotic triangle in the context of taxonomic reference.

## Taxonomic reference modeling and the semiotic triangle

The semiotic triangle, or triangle of meaning/reference, was expounded in Ogden and Richards (1923, though see also Peirce 1998). This influential model of meaning was recently applied to nomenclatural and taxonomic use cases (Remsen 2015, Witteveen 2015a, though see also Jones 1974, Payne 2012). We use the model here with reservations related both to adequacy and completeness. In particular, in its original configuration (figure 1A) the triangle omits the identities and interactions of multiple human (and/or machine) communicators separated in space and time. Overall the model is too simple to encode ‘what concepts are’, or ‘what taxa are’ (Schulz et al. 2008, Franz and Thau 2010, Littlejohn and Foss 2010). In spite of these and other caveats described below, the semiotic triangle is adequate for explaining why computers struggle to match human abilities for differentiating between different kinds of name-facilitated reference services.

**Figure 1.**
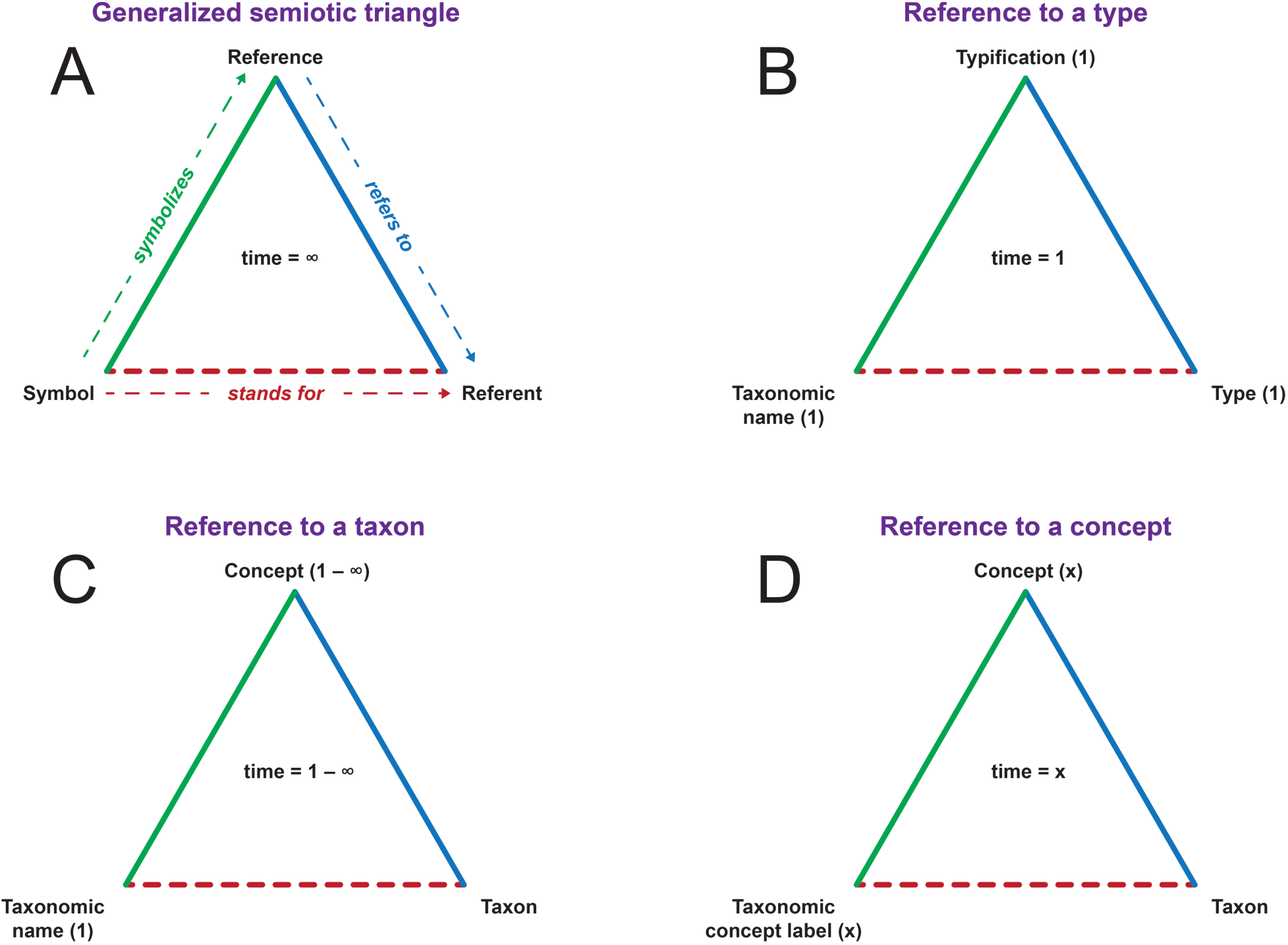
*Taxonomic reference and the semiotic triangle (not incorporating transition). The relation of the symbol and the reference is imputed (as opposed to causal). (A) Generalized semiotic triangle of Ogden and Richards (1923), without phase specification (time = ∞). (B) Reference to a nomenclatural type. A taxonomic name symbolizes reference to (or: stands for) a nomenclatural type at time = 1 when the name is coined and typified. (C) Reference to a taxon. A name coined and typified at time = 1 symbolizes reference to (or: stands for) a taxon at any and all times (time = 1 – ∞). (D) Reference to a taxonomic concept. A taxonomic concept label used at a particular time = x symbolizes the corresponding taxonomic concept at time = x, which in turn refers to the taxon.*

In its generalized version (figure 1A), the triangle has three elements – symbol (also called sign), reference (also called concept or thought), and referent (also called [real world] object). The three resulting relations among paired elements are as follows (Ogden and Richards 1923: 10–12). (1) The symbol *symbolizes* the reference via a causal relation, i.e., via rigid designation mediated by an initial event of baptism and a succeeding causal chain of reference, the latter spanning from that past event to the present (Stanford and Kitcher 2000). (2) The reference *refers to* the referent, again, through a causal relation. (3) The symbol *stands for* the referent in an ascribed (“imputed”), indirect relation drawn with dashed (as opposed to solid) lines.

The generalized semiotic triangle can be customized to correspond to each of the three aforementioned kinds of reference services provided by taxonomic names (figures 1B–D). In doing so we begin to model temporal transitions and multi-speaker elements, as appropriate. We will further describe these in subsequent models (figures 2–4).

**Figure 2.**
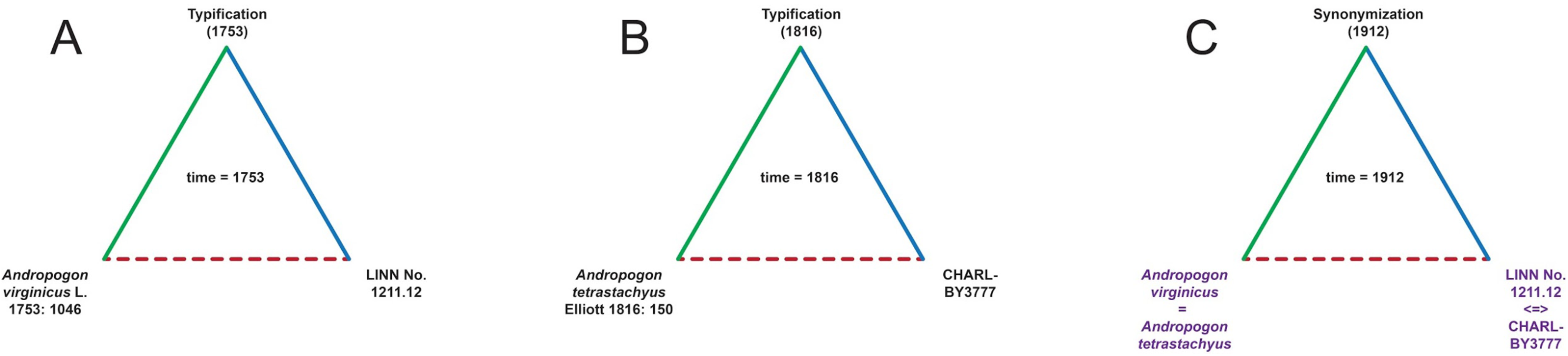
*Transitional model for reference to nomenclatural types (expansion of figure 1B). (A) Reference to the type specimen that typifies the name* Andropogon virginicus *at time = 1753. (B) Reference to the type specimen that typifies the name* Andropogon tetrastachyus *at time = 1816. (C) Act of heterotypic synonymization of the type specimens and corresponding names, where the name* Andropogon virginicus *has Priority.*

**Figure 3.**
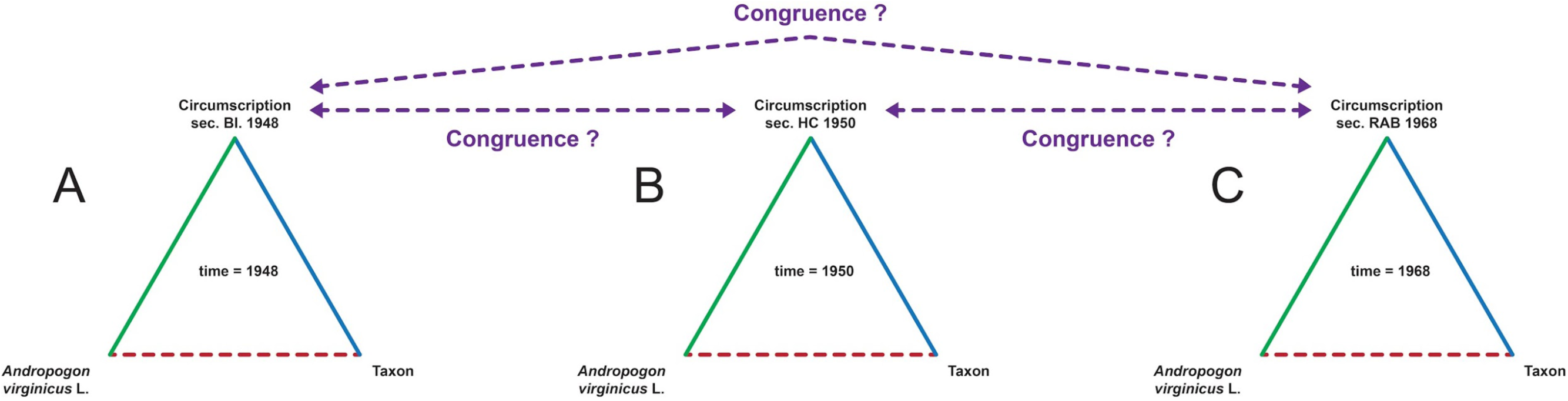
*Transitional model for reference to taxa (expansion of figure 1C). The models (A), (B), and (C) reuse the name* Andropogon virginicus *while symbolizing three temporally distinct phases of taxonomic concept circumscriptions. The semantic congruence across the taxonomic concepts is uncertain (purple lines), and the symbol (name) in effect symbolizes the entire taxonomic concept lineage (1948 to 1950 to 1968). The extent to which each circumscription refers to the taxon is not specifiable (beyond the identity of the type specimen).*

**Figure 4.**
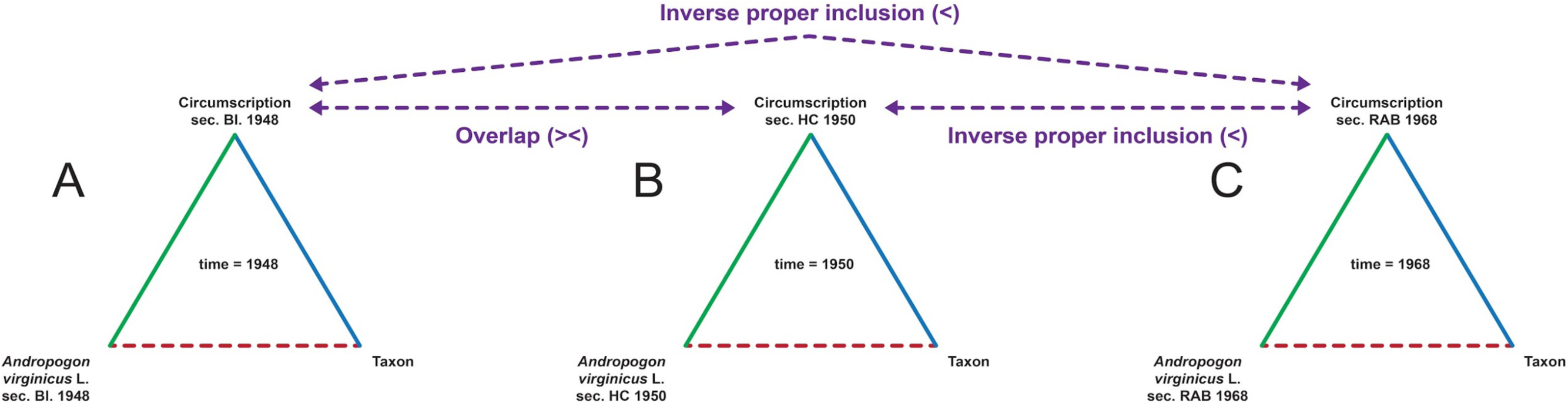
*Transition model for reference to taxonomic concepts (expansion of figure 1D). Each of the models (A), (B), and (C) is uniquely identified with the taxonomic concept label that symbolizes the contextualized taxonomic concept circumscription of 1948, 1950, or 1968, respectively. An additional semantic layer (purple lines) provides logic-amenable taxonomic provenance information, in this case utilizing RCC-5 relationships.*

In the model in which taxonomic names refer to types (figure 1B), the symbol is the taxonomic name as used at the time (t = 1) of the act of typification. That time-stamped, Priority-granting act plays the role of the reference, and the type (specimen) is therefore the referent. In our example (figure 2A), the name *Andropogon virginicus* L. 1753: 1046 (symbol) is typified by (reference) the type LINN No. 1211.12 (referent).

We can immediately observe that the model is problematic. For instance, not all types in the Linnaean system are specimens – names can act as types too (yet are not objects in an immediate, physical sense). Moreover, if names do indeed refer to their types by necessity (Witteveen 2015a, 2015b), then this relation of symbol to referent is causal, not (just) ascribed. We mention in passing that typification of higher-level taxonomic names (beyond the family rank) is not enforced by the *Codes* of nomenclature (Dubois 2015). Hence the nomenclatural model (figure 1B) does not hold true for these names. Lastly, typification is an act of creating meaning relations ostensively, i.e., by pointing to an object (Stanford and Kitcher 2000). The triangle is arguably more appropriate for accommodating reference practices that include descriptivist, intensional components.

The model where names refer to taxa also utilizes the taxonomic name in the role of the symbol (figure 1C). Typically, this is the valid name carrying Priority, or having been conserved in light of stability considerations (Vences et al. 2013). It is often the same name (identifier string) used to refer to the nomenclatural type. One significant feature of this model is the underspecified relationship of the symbol to the reference, i.e., the taxonomic concept. Recall that the taxonomic name, as typified at t = 1, functionally stands for the taxon in this model. In other words, the taxonomic concept is held stable *so that* the name-referent relation can be established with the desired degree of semantic precision. Taken in isolation, the model is not suited to represent the individualized taxonomic provenance of the reference, instead providing an atemporal or provenance-unaware view of the reference (hence the annotation t = 1 – ∞). In the minds of human speakers, the model can nevertheless succeed in communicating about biological properties of the taxon for which the name stands.

The model that emphasizes the relation of names symbolizing concepts is similar to the preceding one in terms of having an intensional, descriptivist focus; however it is temporally specified and therefore provenance-aware (figure 1D). The symbol reflects this temporal dimension by being individuated *according to* the human authors publishing the corresponding taxonomic concept (Berendsohn 1995). Examples of such symbol individuation include: (1) *Andropogon virginicus* L. sec. Hitchcock and Chase (1950) and (2) *Andropogon virginicus* L. sec. Radford, Ahles and Bell (1968). We refer to the temporally individuated symbols as taxonomic concept labels to differentiate them from the taxonomic names that act as symbols in the preceding models (figures 1B and 1C). Notice however that every taxonomic concept label may also contain a type-anchored taxonomic name. This means that taxonomic concepts labels are generally suited to function in the nomenclatural and taxonomic reference domains.

The taxonomic concept label precisely symbolizes one particular reference. Both the symbol and the reference carry the time stamp t = x (where x is 1950, or 1968, in the above examples). While this precise individuation of the concept does not preclude the latter from referring to the taxon, it does so explicitly in relation to the taxonomic identity specifiers of the reference under consideration. The symbol stands for the taxon in this reference-specific sense.

## Transitional taxonomic reference models

Communication via taxonomic names is intended to succeed over increasing spatial scales and across generations of communicators. Often the referential contexts in which names were originally applied are modified through subsequent usages. We therefore ask: how can each model (figures 1B–1D) be expanded to account for possible temporal transitions? Another way to frame this question is to investigate the precision with which the three elements of the semiotic triangle (figure 1A) are identified and semantically linked to each other across multiple temporally succeeding semiotic triangles, as shown in figures 2–4.

In the example shown (figure 2), the symbol *Andropogon virginicus* L. typifies the referent type specimen LINN No. 1211.12 at t = 1753 (figure 2A). Subsequently, the symbol *Andropogon tetrastachyus* Elliott typifies the referent type specimen CHARL-BY3777 at t = 1816 (figure 2B). Then at t = 1912, an act of heterotypic synonymization occurs (figure 2C). We note that the semiotic triangle may be increasingly inappropriate, or at least unnecessary, to represent this act. Regardless, the output of the transition model is readily characterized. The two symbols are now considered nomenclaturally synonymous (annotated with “=”). The corresponding type specimens, while still only anchoring their respective names (figures 2A and 2B), are closely and ostensively linked referents (“<=>”). Thus the nomenclatural transition model clearly shows provenance awareness in reference to nomenclatural types. We have to defer more detailed analyses of the semantics of Code-mandated (type) relationships for future expositions.

Our comparison of the other two transition models utilizes parts of the *Andropogon* use case as developed in Franz et al. (2015b). In the first of these, the same symbol (identifier string) *Andropogon virginicus* L. is used at t = 1948, t = 1950, and t = 1968 (figure 3). In each phase of the model, the symbol effectively symbolizes the corresponding taxonomic concept, i.e., the intensionally formulated taxonomic diagnosis of one of the three published accounts. The extent to which the three references are semantically congruent – beyond their sameness in the nomenclatural domain – is underspecified. The syntactic constraints on the symbol are such that it is provenance-unaware *by design* (figure 2C). This means that the symbol is generally well suited to symbolize the entire, potentially open-ended transition sequence of references at once, i.e., to symbolize the taxonomic concept *lineage.* But it is not suited to identify any specific phase in the temporally evolving model. This also means that the relative congruence across temporally succeeding references (taxonomic concepts) and referents (taxa) cannot be assessed under this model.

Relaxing the syntactic constraint (same identifier string) on the symbol allows exactly one provenance-aware taxonomic concept label to appear in each semiotic triangle in the transition sequence (figure 4). The respective reference is thereby also uniquely symbolized. The ability to identify the entire transition model – the taxonomic concept lineage – with an individual symbol is lost. To do so, multiple symbols would have to be utilized and linked (three in this example). This transitional representation is capable of assessing the extent to which the constituent references (and therefore taxa) are reciprocally semantically congruent, or not. It turns out (figure 5) that the concepts *A. virginicus* L. sec. Blomquist (1948) (figure 4A) and *A. virginicus* L. sec. Hitchcock and Chase (1948) (figure 4B) overlap intensionally, and each is properly included in *A. virginicus* L. sec. Radford, Ahles and Bell (1968) (figure 4C).

**Figure 5.**
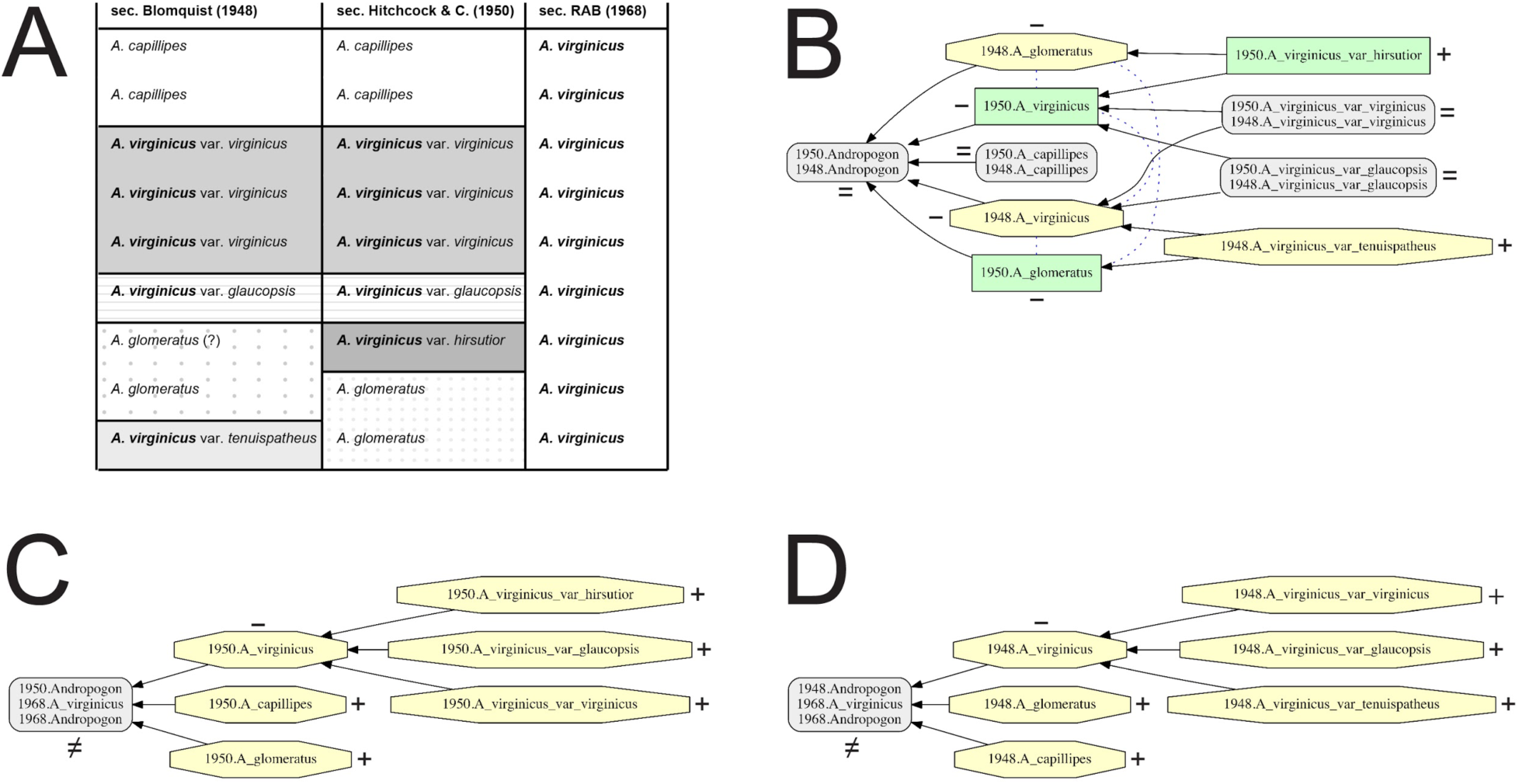
*Logic-amenable representations of taxonomic concept provenance for the name* Andropogon virginicus *participating in the three taxonomic concept labels* Andropogon virginicus *sec. Blomquist (1948),* Andropogon virginicus *sec. Hitchcock and Chase (1950), and* Andropogon virginicus *sec. Radford, Ahles and Bell (1968). Adopted from Franz et al. (2015b). (A) Tabular alignment of lowest-level taxonomic concepts; horizontal row identity indicates taxonomic concept congruence. Taxonomic concept labels that entail the name* Andropogon virginicus *are rendered in bold font. (B) Pairwise alignment visualization of the classifications of Hitchcock and Chase (1950) and Blomquist (1948), as generated with the Euler/X software toolkit. Taxonomic concept labels are abbreviated. Grey rectangles with rounded corners indicate congruent concept regions; green rectangles with sharp corners indicate concepts unique to the later taxonomy; and yellow octagons indicate concepts unique to the earlier taxonomy. Solid lines indicate proper inclusion, whereas blue dashed lines indicate overlap. Additional annotations: = : congruent concepts, identical names; ≠: congruent concepts, non-identical names; + : unique concept, unique name; – : unique concept, non-unique name. (C) Alignment of the classifications of Radford, Ahles and Bell (1968) and Hitchcock and Chase (1950). (D) Alignment of the classifications of Radford, Ahles and Bell (1968) and Blomquist (1948). See Franz et al. (2015a, 2015b) for additional explanation.*

## Transitional reference framing: human cognitive abilities and logic constraints

We may now diagnose why computational logic struggles to accommodate the kinds of reference services and successes that humans have come to expect when using taxonomic names. The cause for the struggle is diagnosable as an instance of the frame problem (McCarthy and Hayes 1969, Shanahan 2009), where knowledge of relevant yet not explicitly stated conditions is necessary in order to reason intelligently.

In nomenclaturally oriented publications, is it commonplace for authors to shift from one kind of temporally evolving reference model to the next (figures 2–4), albeit using primarily or exclusively taxonomic names as symbols. In countless other publications in comparative biology where nomenclatural issues are not the central interest (including on-line, networked database environments), human authors will nevertheless shift abundantly and seamlessly among the two models that entail taxonomic concepts (figures 3–4). Both kinds of reference services – reference to concepts and to taxa – are essential components of effective communication in this context. For instance, a comparative phylogenetic study may commence with introductory and method sections where previously published taxonomic concepts are symbolized to set the stage for the new analysis. Once the phylogenetic inference is completed and symbols are (re-) assigned accordingly (in the results and discussion sections), the symbols now stand for the corresponding taxa whose phylogenetic properties are under examination. Indeed, shifting back and forth between the two models (figures 1C and 1D) may occur very frequently in any section of the publication because both the history of human taxonomic theorizing and the properties of biological taxa are relevant. In many cases the varying models are applied without explicitly and consistently individuating taxonomic concepts labels as symbols.

To review – human speakers routinely shift between concept- and taxon-symbolizing, semantically evolving reference models in their comparative biological publications. They do so while using primarily or exclusively taxonomic names as symbols. We suggest here that this practice is acceptable, and may even been preferable from their perspective, because of shared and uniquely human abilities to contextualize symbols that nevertheless remain structurally underspecified (symbolizing entire, open-ended lineages of taxonomy concepts) from the viewpoint of computational logic. This means that humans must in effect and implicitly generate more, and more semantically granular, symbols (identifiers) for mutually perceived concepts and taxa than their publications explicitly contain. Moreover, these implicitly communicated, more granularly individuated identifiers are essential for explaining the success of human-to-human communications within and across reference models and services.

When two similarly trained botanists residing in the southeastern United States and living today (2015) communicate about *Andropogon virginicus,* there is a good likelihood that each can reliably navigate across all semantic elements and relations displayed in figures 2–5. This is part of their shared ability to solve the frame problem in this example. Quite likely both will have formed additional implicit symbols and relations for concepts represented in more recently published accounts that utilize the name *Andropogon virginicus* than that of Radford, Ahles and Bell (1983) (Franz et al. 2015b). Neither might intuitively default to the taxonomic concepts as circumscribed in 1753 when the name *Andropogon virginicus* was typified. Such 18th or 19th century concepts are inter-subjectively ‘known’ to be largely out of use. In other words, the human communicators have rich and time-tested mental abilities and communication repertoires at their disposal to implicitly supplement the proper frame conditions and added degrees of symbol granularity, and thereby succeed in understanding each other’s name usages and concepts and their provenance.

From the perspective of computational logic, in turn, humans are carrying out semantically diverse suites of reference services while using taxonomic names that appear syntactically incapable of facilitating these services. Without additional, implicitly accessed knowledge of context, computers are not able to represent any of the semiotic triangles displayed in figures 4–5 individually, or infer the provenance among their elements.

In asking computers to represent and reason over taxonomic (provenance) knowledge via taxonomic names in the ways we are accustomed to, humans are asking computers to perform reasoning services for which no logically adequate syntactic and semantic structures are provided. Unless the semantic relations across all phases in the transition model are indeed stable and congruent, the services cannot reliably materialize.

## Why is there not enough semiotic structure for computers?

How to move forward from this diagnosis? Our intention is not to convince humans to neglect their inherent and acquired talents for contextualizing taxonomic symbols and concepts when communicating with each other. But we might inquire why an only implicitly provenance-aware system for identifying and integrating taxonomic knowledge came to such wide acceptance among humans (Schuh 2003, Dayrat 2010). Outlining the answers might produce additional insights into the design needs for logic-facilitated taxonomic reference services.

The aforementioned constraints on symbols are enforced through the Codes of nomenclature. It is therefore important to examine why the semiotic conventions represented in the widely accepted Codes are they way they are. Here we can only sketch out pathways towards more comprehensive answers. For instance, we note that an overarching goal of the Codes is to promote long-term stability in naming. This is achieved, first and foremost, by the Principle of Priority, which mandates that the oldest available name for a “taxon” (in our representation – taxonomic concept lineage, figure 4) is the valid one. Under certain conditions the Principle can be overridden, and instead another name is conserved as valid for a taxon. Typification is required at the lower taxonomic name ranks to which the Codes apply, and type identity is the primary mediator of reference identity.

Immediately we observe discrepancies between Code-compliant symbolizing and (e.g.) the logically tractable reference models displayed in figures 1D and 4. The Codes explicitly enforce the use of type-anchored names, not of taxonomic concept labels. Reusing the same valid name in publications that succeed the original, type-anchoring treatment is not only encouraged but required, as long as the corresponding type is entailed in the subsequent taxonomic concept of the transition model. Furthermore, the Codes do not prescribe specific and consistent methods to model taxonomic concept provenance. Although the reference (circumscription) and referent (type, taxon) are separately recognized in many sections of the Codes, the provenance of name usages (homonymy, synonymy) is generally established in relation to the identity of nomenclatural types (figures 1B and 2). Indeed, the Codes emphasize that nomenclatural conventions should not “infringe upon taxonomic judgment” (e.g., ICZN 1999). By freely permitting that judgment to establish itself and evolve ‘on top of’ regulated notions of nomenclatural (type-centric) provenance, the Codes remain silent on whether and how to achieve to the complementary taxonomic provenance semiotics that computational logic would require to match human abilities for contextualizing name usages.

We may conjecture that many hard-wired features of the Linnaean reference system are optimized (or nearly so) for human-to-human communication and learning about nature’s diversity and hierarchy. Notable human cognition-optimized features include the following. (1) The mandated reuse of names symbolizing taxonomic concept lineages in effect limits the number of different name strings that humans have to cognitively manage. (2) The use of binomial names for species concepts mirrors the Aristotelian method of creating genus-differentia definitions (Ereshefsky 2001). (3) The use of standardized endings for certain taxonomic name ranks in hierarchical classifications, typically limited to less than 5-15 ranks and endings, is consistent with human classificatory habits as observed in diverse folk biologies (called “cognitive universals” in Atran 1998, see also Berlin 1992). (4) The requirement of combining intensional (descriptivist, feature-based) and ostensive (causal, member-based) elements in taxonomic concept circumscriptions (references) is consistent with the hybrid (causal/descriptive) theory of reference and may be uniquely suited for human cognitive processing (Evans 1982, Devitt and Sterelny 1999, Franz and Thau 2010). (5) Single-specimen typification is both an implementation of the causal theory of reference (Witteveen 2015a, 2015b) – thereby ameliorating misunderstandings due to inadequate (partial, erroneous) description – and an act whose sensible interpretation requires cognitive abstraction from the exemplar to the taxonomic concept (Winsor 2003). (6) The lack of jurisdiction of the Codes over higher taxonomic levels (order level and above) suggests that intensional reference practices predominate over type-based definitions at these larger taxonomic scales where circumscriptions by ostension would become increasingly data-intensive (Franz and Thau 2010). In brief, we suggest that many semiotic properties of the Linnaean reference system are not arbitrary in their fundamental specification, and instead are closely aligned with cognitive and communicative preferences of humans in particular. Furthermore, the system’s optimizations benefitting human communication frequently work to the detriment of modeling contextualized reference services in computational logic.

Computational logic is not immediately constrained by, or optimized for, adherence to human cognitive universals in the context of framing taxonomic communication (see points 1–6 above). Hypothetically speaking, if the objective is to logically represent the transition model of figure 4, computers have no need at all to reuse symbols for references authored at separate times (1). Moreover, the symbols would not need to be mono- or binomial, or of limited length and thereby easily pronounceable and memorable for humans (2). Rank endings would not have to be inherently embedded in the symbols, as this information could be allocated in the associated reference (3). Instead it would be perfectly suitable in the ‘eyes’ of computational representation to assign (e.g.) globally unique 128-bit strings for every taxonomic concept label. Handling trillions of such unique symbols for every reference (taxonomic concepts) would be well within reach. The provenance of uniquely symbolized concepts could be computed over an additional layer of logically actionable reference-to-reference linkages. Each linkage might be based on potentially large sets of diverse data types, including explicitly assigned specimens (type or otherwise), entailed lower-level concepts (if higher-level concepts are symbolized), specified phenotypic and genotypic traits, and other semantics that measure taxonomic concept congruence, or the lack thereof (4, 5). The system should scale from the lowest to the highest taxonomic concept levels without needing to shift weights between intensional and ostensive circumscriptions due to increasing data volumes and other cognitive constraints (6). The system could be queried for taxonomic concept identity and provenance information and reliably represent the contextuality and semantic linkages of every symbol and semantic element of the corresponding reference(s) (figure 4).

## Can we attain the best of both worlds?

The above, hypothetical solution of using highly contextualized taxonomic concept identifiers and linkages may approximate optimality for computers. But it does so at the expense of rendering human communication about taxonomic symbols and references and their provenance virtually impossible. Hence we might ask: what would an attainable and desirable middle ground solution look like? Is the ‘best of both worlds’ – i.e., taxonomic symbols that are maximally aligned with human cognitive capacities yet which are also precisely framed for logic-based representation and reasoning – an option?

We think so. In fact, Berendsohn’s (1995) proposal to consistently individuate taxonomic concept labels represents one major innovation in transitioning to the reference services displayed in figure 4. This is achievable without losing any of the Linnaean system’s inherent optimizations vis-à-vis human communication. Several taxonomic publications or software projects have adopted the more granular symbology and proceeded to build layers for computing provenance between taxonomic concepts on top of the contextualized name usages (e.g., Koperski et al. 2000, Pullan et al. 2000, 2005, Cui 2012, Franz et al. 2015a, Jansen and Franz 2015). Avibase (Lepage et al. 2014) is one of the most thorough and impacting realizations of this semiotic approach. This database environment manages to uniquely symbolize 844,000 species-level and 705,000 subspecies-level taxonomic concepts spanning across 151 checklists of birds published over 125 years. Provenance between references is defined through an integrated layer of Region Connection Calculus (RCC-5) relationships (Franz and Peet 2009). The added semantics facilitate the recognition of 38,755 taxonomically distinct concept lineages, each with one to many references. They furthermore show that only 11 of 19,260 (∼ 1 in 1750) taxonomic name/concept combinations are both syntactically and semantically unique; thus having the symbol:reference cardinality of 1:1 throughout the entire taxonomic reference and provenance environment. All other combinations are in need of framing to become uniquely identifiable.

It is perhaps not accidental that Avibase (Lepage et al. 2014) has acquired such highly contextualized symbology and taxonomic concept provenance semantics when most other comparative biology data environments continue to adhere to reference practices that are less computationally accessible. Avibase focuses primarily on representing classifications below the family-level name rank. It must accommodate one or more new taxonomies each year, published as updated checklists by different global or regional authorities such as the American Ornithologists’ Union or the International Ornithological Committee. Each checklist synthetically annotates an authority’s time-stamped taxonomic perspective using fairly homogenous and simple semiotics. Neither featured-based circumscriptions nor specimens are represented. There is immense use of these taxonomies, whose frequency of publication and complexity of semantic relations rapidly exceeds human cognitive abilities for reliable reconciliation. It is as if for once humans were ‘put in the shoes of computers’, and realized that the desired taxonomic reference services at Avibase’s scale and density of transition could only be obtained through more structured semiotics.

## What about taxa?

The taxon (referent) element is kept purposefully vague in our reference models with regards to temporal specifiers and in comparison to the symbol and reference elements (figures 1C, 1D, 3 and 4). We hope that the models can thereby support at least two necessary readings. Under the first of these, the taxon’s identity is considered stable across the temporally succeeding semiotic triangles. The taxon, roughly, is represented as an evolutionary unit – acknowledging many ontological complexities this might entail (Boyd 1999, Stanford and Kitcher 2000, Ereshefsky 2001, Schulz et al. 2008, Brigandt 2009) – whose identity is not relevantly altered in the course of less than three centuries of (post-)Linnaean classifying. This reading can account for any particular taxonomic concepts to variously succeed, or fail, in referring to the evolutionary identity of the taxon. The history of taxonomy can thus be viewed as one in which the intensionalities of published concepts successively (if not linearly) approximate the identities of taxa. In other words, the transition models of figures 3 and 4 support the view that taxa have an ontological standing, which the practice and gradual advancement of taxonomic research and classifying can successfully strive to uncover, at some level of precision. Taxa ‘are what they are’, and taxonomy is becoming increasingly better at understanding their identity.

The second reading allows the identity (identities) of the taxon (taxa) to vary across multiple semiotic triangles. Here the identity of the taxon is taken as more contingent on the classifiers’ intensionality. For instance, the two taxonomies displayed in figures 5B and 5C, or in figures 5C and 5D, are incongruent mainly by virtue of the 1968 taxonomy’s coarse granularity. The corresponding authors may well understand why alternative classifications recognize more granular taxonomic concepts. But they might not judge the differences between these lower-level concepts sufficiently significant to be mirrored in taxonomic name resolution. Meanwhile, the more finely resolved referents could have causally aligned referents in nature. In that case the corresponding taxa are non-identical, and the lack of congruence is rooted in the interplay of causal phenomena and differential taxonomic assessments of the relevance of these phenomena in the context of comparative biological classification. Under this reading, then, taxa are partly ‘made’ by taxonomists, in the sense that different theory-driven taxonomic weighting schemes may result in the recognition of more, or less, granular concepts, or incongruent integrations of such concepts into higher-level concepts, where each alternative reference scheme remains alignable with causally sustained entities in nature.

In summary, the transitional taxonomic reference models (figures 3 and 4) accommodate complementary readings of the human history of taxonomy creation and refinement. Under one interpretation, reference to (relatively) stable evolutionary taxa can successively approximate high measures of reliability. Under another interpretation, alternative theory-informed conceptualizations can weigh evolutionary phenomena differentially but nevertheless remain empirically adequate. Under either reading it is necessary to represent taxonomic concept provenance as outlined above to leverage computational logic and services.

## Metawork: taxonomic reference services for comparative biological data environments

Comparative, taxonomically referenced biological data are becoming increasingly digitized, networked, redistributed, and reutilized in syntheses and analyses and that expand or alter their original context of generation. The era of the mostly analog, and perhaps aspirationally definitive taxonomic knowledge product – with well-defined boundaries of context, authorship, and publication – might gradually come to an end. In particular, comparative biologists are now developing virtual environments that are both synthetic with respect to existing knowledge, and open to continuous integration of new empirical information (e.g., Lepage et al. 2014, Deans et al. 2015, Smith et al. 2015). In developing these open-ended platforms, they must integrate sets of taxonomic concepts originating from different source publications. Often the taxonomic scope of each source is thereby complemented and modified in ways that the original work could not have anticipated. Once the initial phase of assembling multi-sourced external accounts is completed, the environments effectively propagate new theoretical content (Leonelli 2013). Over time the content evolves, resulting in new phases of the transitioning environment that may empirically conflict with previously endorsed content.

Enabling broad and deep analyses of taxonomic diversity and relationships is perhaps the most explicit goal of comparative, synthetic information environments. This means that, at any given time, taxonomic names used in the environments are intended to serve the function of *standing for* taxa (figure 1C) whose biological properties are comparatively analyzed. However, our models indicate that without providing additional symbol resolution in the form of taxonomic concept labels (figure 1D), the information environments cannot achieve taxonomic provenance awareness.

Virtual comparative information environments that are open to accommodating taxonomic advancements face new syntactic and semantic design opportunities. We are of the view that environments that specialize in representing the legacy of human taxonomy making should explore the conventions displayed in figures 4–5. But what about more generally positioned platforms – can these other systems do well and evolve without providing logic-amenable taxonomic reference services? With reservations, we suggest that the same logic constraints apply (Franz and Thau 2010). This means that even if an information environment’s primary focus is to facilitate comprehensive taxonomic/phylogenetic syntheses and/or contingent evolutionary analyses, the environment will nevertheless require adequate modeling of taxonomic concept identity and provenance as displayed in figure 4. Such provenance modeling is needed to ensure that open-ended environments can continuously expand in scope while retaining the ability to identify and logically integrate their internal, transitioning taxonomic semiotics.

Adding computationally actionable annotation and integration layers can be considered metawork, i.e., the work of organizing work (Gerson 2008, 2009, Brigandt 2010). Taxonomic information systems that have benefitted from such metawork are logic-enabled. They can therefore provide all relevant taxonomic reference and provenance services to taxonomic experts, taxonomy users, and – critically – other similarly empowered computational systems. Designing, implementing, and maintaining these logic-based systems will facilitate rationalized coordination of transitioning taxonomy environments, benefitting both humans and machines. Absent such metawork, the costs of investing into the content of open-ended comparative biological information environments may become unsustainable.

## Caveats and future directions

In this paper we explore numerous interconnected themes in deliberately broad strokes. We recognize that many representations and linkages need further, detailed analysis. For instance, throughout we have been largely silent on the semantic content of taxonomic concepts, and how that content might be logically reconciled across accounts (e.g., Franz and Peet 2009, Cui 2012, Franz et al. 2015a). Understanding the extent to which logic representations of rich taxonomic semantics are possible is nevertheless critical for judging the benefits of these representations over conventional, nomenclatural provenance. The challenge is relevant both to representing the legacy of human taxonomy making and moving towards future, more logic-amenable, reference practices. In many cases humans and computers will have to accept uncertainty in reconciling taxonomic concepts.

The conditions and significance of partial failure in taxonomic reference in human-to-human communication are also worth deeper analysis. In particular, we do not mean to suggest that humans always perform well in contextualizing the taxonomic names in the figure 3 model, or that no computationally effective representations of taxonomic provenance are possible unless one implements exactly the figure 4 model. We also point out that for many comparative biological analyses, the taxonomic provenance resolution afforded by taxonomic names is adequate to support the therein-intended inferences, and likely many other related inferences.

The issues arise when the contexts of symbolizing taxa expand greatly over time. Our central argument is that the conventional practices of identifying and reconciling taxonomic concepts are systemically, even justifiably, biased to favor human cognitive strengths and deemphasize our cognitive weaknesses in comparison to computers. Hence the Linnaean system allows *us* to achieve reliable taxonomic reference services, albeit at human cognition-limited volumes. For deeper temporal scales and greater data volumes the prospects are less encouraging (Franz et al. 2008, Lepage et al. 2014). If we expect comparative biological data environments to scale up concept reference services to match the full body of taxonomic knowledge, then we must improve the syntax and semantics of human taxonomy making for computers.

## Acknowledgments

The authors are grateful to Hong Cui, Bertram Ludäscher, and Jonathan Rees for helpful feedback on this subject. Support of the authors’ research through the National Science Foundation is kindly acknowledged (NMF: DEB–1155984, DBI–1342595; BWS: SES– 1153114).

